# A mitochondrial insertion directs substrate selection and engagement by ClpX

**DOI:** 10.64898/2026.02.23.707539

**Authors:** Adam DeCosta, Rebecca Barrick, Julia R. Kardon

## Abstract

The force-generating AAA+ ATPase domain of protein unfoldases is specified for many substrates and other functional partners through elaboration with accessory domains. Mitochondrial homologs of the unfoldase ClpX contain an insertion within the AAA+ domain that is absent in bacterial homologs. This mitochondrial insertion (MI) maps to the substrate-encountering face of ClpX, leading us to hypothesize that the MI directs interactions of ClpX with mitochondrial substrates, the best-characterized of which is the first enzyme in heme biosynthesis, ALAS. We find that the MI is critical for both recruitment and activation of ALAS by *S. cerevisiae* ClpX. The MI was dispensable for heme-induced, adaptor-mediated degradation of ALAS by human CLPXP, but contributed to adaptor-independent recruitment of the model substrate casein for degradation. Although truncation of the MI moderately perturbed ATPase activity in both yeast and human ClpX, this effect could be uncoupled from the requirement for the MI in the efficiency of ALAS activation by targeted mutagenesis. The MI therefore can serve both to recruit a substrate to mitochondrial ClpX and to accelerate its processing by the AAA+ motor.

## INTRODUCTION

AAA+ family unfoldases provide essential control of protein biogenesis, repair, regulation, and quality control^1^. These enzymes share a highly conserved AAA+ ATPase domain that grips protein substrates to apply the mechanical force that drives protein unfolding^2^. This core mechanism must be controlled and directed for specific substrates and functions. Intensive study of several bacterial AAA+ unfoldases has demonstrated that like several other well-studied members of the larger AAA+ family, accessory domains are responsible for directing the AAA+ domain to specific functions. For AAA+ unfoldases, these accessory domains directly mediate specific interactions with substrates or with adaptor proteins and other functional partners^3^. Mitochondrial AAA+ unfoldases are closely related to those of their bacterial relatives, but in many cases these accessory domains have diverged widely between bacterial and mitochondrial homologs, for which substrate repertoires and recruitment strategies are much less characterized.

One example of such divergence is the AAA+ unfoldase ClpX, which is widely conserved in bacteria and as a mitochondrial protein in eukaryotes. In both bacteria and mitochondria, ClpX can act to regulate substrates on its own by remodeling them^4,5^ and by unfolding coupled to degradation in complex with its partner peptidase ClpP^6^. ClpX is essential for viability in many bacteria^7^, its deletion is early-embryonic lethal in mice^8,9^, and mutations in ClpX or ClpP underly two mitochondrial syndromes^10^, underscoring the broad importance of the regulation of bacterial and mitochondrial proteomes by ClpX. Proteomic, cell biological, and biochemical data suggest that the substrate repertoire of mitochondrial ClpX homologs has diverged widely from known bacterial substrates^5,11,12^. A broadly conserved function of bacterial ClpX, the degradation of stalled translation products directed by the ssrA tag and the adaptor protein SspB, appears to have been lost in the evolution of mitochondria, which lack genes encoding SspB, and with few exceptions, the ssrA tag^13^. Although both bacterial and mitochondrial ClpX homologs recruit adaptor proteins by a related N-terminal zinc-finger domain, the single demonstrated adaptor protein for mitochondrial ClpX^14^ appears to be restricted to metazoans, and no bacterial adaptors appear conserved as mitochondrial proteins. Mitochondrial ClpX homologs also contain an element of ∼30-70 amino acids in length that is absent in bacterial homologs. This mitochondrial insertion (MI) is inserted between two helices of the core AAA+ domain, emerging from the substrate-engaging face of the ClpX hexamer. We therefore hypothesized that the MI could contribute to substrate recruitment by mitochondrial ClpX. Additionally, the MI has been reported to impair oligomerization, ATPase activity, and degradation of the model substrate casein by human CLPXP^15^.

The best-characterized substrate of mitochondrial ClpX is the first enzyme in heme biosynthesis, ALAS. In *S. cerevisiae*, ClpX activates ALAS by partial unfolding to accelerate incorporation or replacement of its cofactor, pyridoxal phosphate (PLP)^5,16^. In metazoans, heme also induces negative-feedback degradation of ALAS by ClpXP^10,14,17^. The adaptor protein POLDIP2 recognizes a heme-bound motif in ALAS and induces degradation of ALAS by ClpXP^14^. Although this appears to be the dominant activity of ClpX toward ALAS in humans, this negative feedback mechanism is absent in *S. cerevisiae*, which lacks both POLDIP2 (which is restricted to metazoans) and ClpP. Here, we establish that the MI is critical for activation of ALAS by *S. cerevisiae* ClpX. The MI is not required for adaptor-mediated degradation of ALAS by human ClpX, but modulates adaptor-independent recognition of the model substrate casein. Our kinetic analysis of ALAS activation indicates that the MI contributes both to recruitment of ALAS and, unexpectedly, to the maximal rate of activation. These data indicate that the MI can direct the activity of ClpX both through substrate selection and by contributing to efficient engagement of substrate by the core AAA+ unfoldase.

## RESULTS

The mitochondrial insertion (MI) is a ubiquitous, small (∼30-70 amino acids) sequence insertion among mtClpX homologs, although its sequence diverges substantially across the eukaryotic phylogeny (Fig. 1A). The MI is predicted to project from the face of the mtClpX hexamer at which substrates are engaged before translocation through the central pore, suggesting that it could contribute to substrate recruitment. In the yeast ClpX homolog, Mcx1, the MI is predicted by AlphaFold3 with low-to-moderate confidence to form a helix-loop-helix structure that bundles with an α helix provided by the small element N-terminal to the AAA+ domain (Fig. 1B). The MI of the human ClpX homolog (CLPX) is predicted with very low confidence as a mixture of helix and coil; although depicted as packing against the N-domain of CLPX, there is no confidence in this conformation (Fig. 1C, S3).

**Figure 1.**
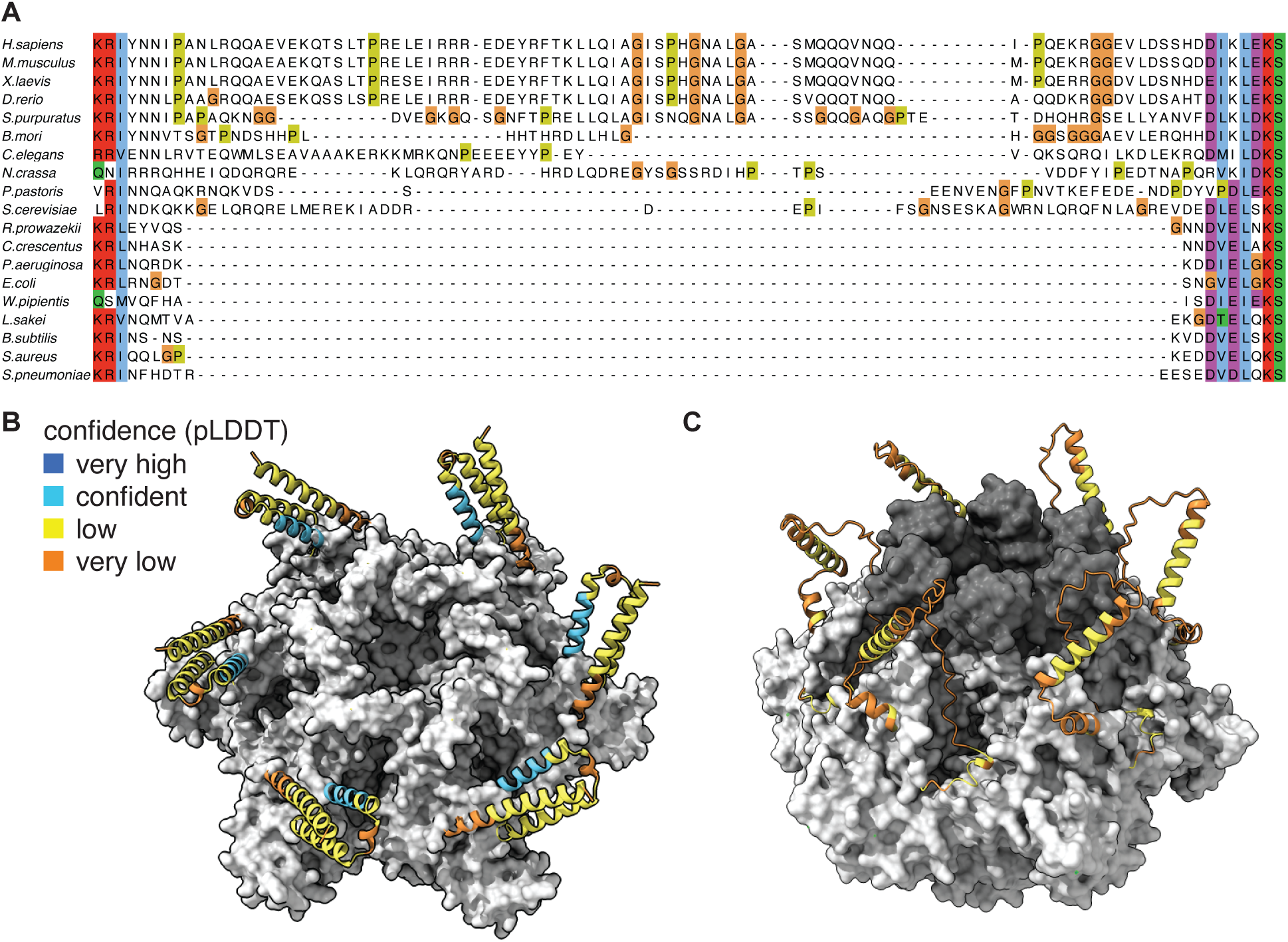
Mitochondrial ClpX homologs contain an element inserted within the AAA+ domain. **(A)** Multiple sequence alignment of bacterial and mitochondrial ClpX homologs at the location of the mitochondrial insertion. Alignment was generated and colored using ClustalOmega with standard settings. Region displayed corresponds to residues 199-287 and 69-134 in human and *S. cerevisiae* ClpX homologs, respectively. **(B)** A model of a hexamer of *S. cerevisiae* Mcx1 generated with AlphaFold3. The AAA+ domain is depicted as a gray surface and ribbon diagrams of the mitochondrial insertion and N-terminal element are colored by AlphaFold3 pLDDT backbone scores (very high, pLDDT >90; confident, 90 > pLDDT >70; low, 70 > pLDDT >50; very low, pLDDT < 50). **(C)** A model of a hexamer of human CLPX generated with AlphaFold3. The AAA+ domain and N-domain are depicted as light and dark gray surfaces, respectively. The ribbon diagram of the mitochondrial insertion is colored by AlphaFold3 pLDDT score as in (B). Very-low-confidence coil at the extreme C-terminus of Mcx1 and N- and C-terminus of CLPX is not displayed.

To investigate the function of the mitochondrial insertion (MI), we first tested whether it contributed to activation of *S. cerevisiae* ALAS. We initially tested this by monitoring the steady-state levels of the product of ALAS, 5-aminolevulinic acid (ALA), in *S. cerevisiae*. ALA levels are reduced in cells lacking ClpX (Mcx1 in yeast) due to reduced ALAS activity^5^. We generated a strain of *S. cerevisiae* in which we replaced the MI with a 3xGly-Ser linker at the genomic locus of Mcx1 (*mcx1^ΔMI^*, 3xGly-Ser replacing residues 81-127 of Mcx1) and then compared ALA levels in this strain to wildtype and *mcx1Δ* yeast. We found that truncation of the MI caused a reduction in cellular ALA equivalent to that observed in *mcx1Δ* cells (Fig. 2A). We detected equivalent levels of Mcx1^WT^ and Mcx1^ΔMI^ protein (Fig. S1), demonstrating that the reduction in ALA we observed in *mcx1^ΔMI^*cells derived from loss of Mcx1 activity rather than reduced protein abundance.

**Figure 2.**
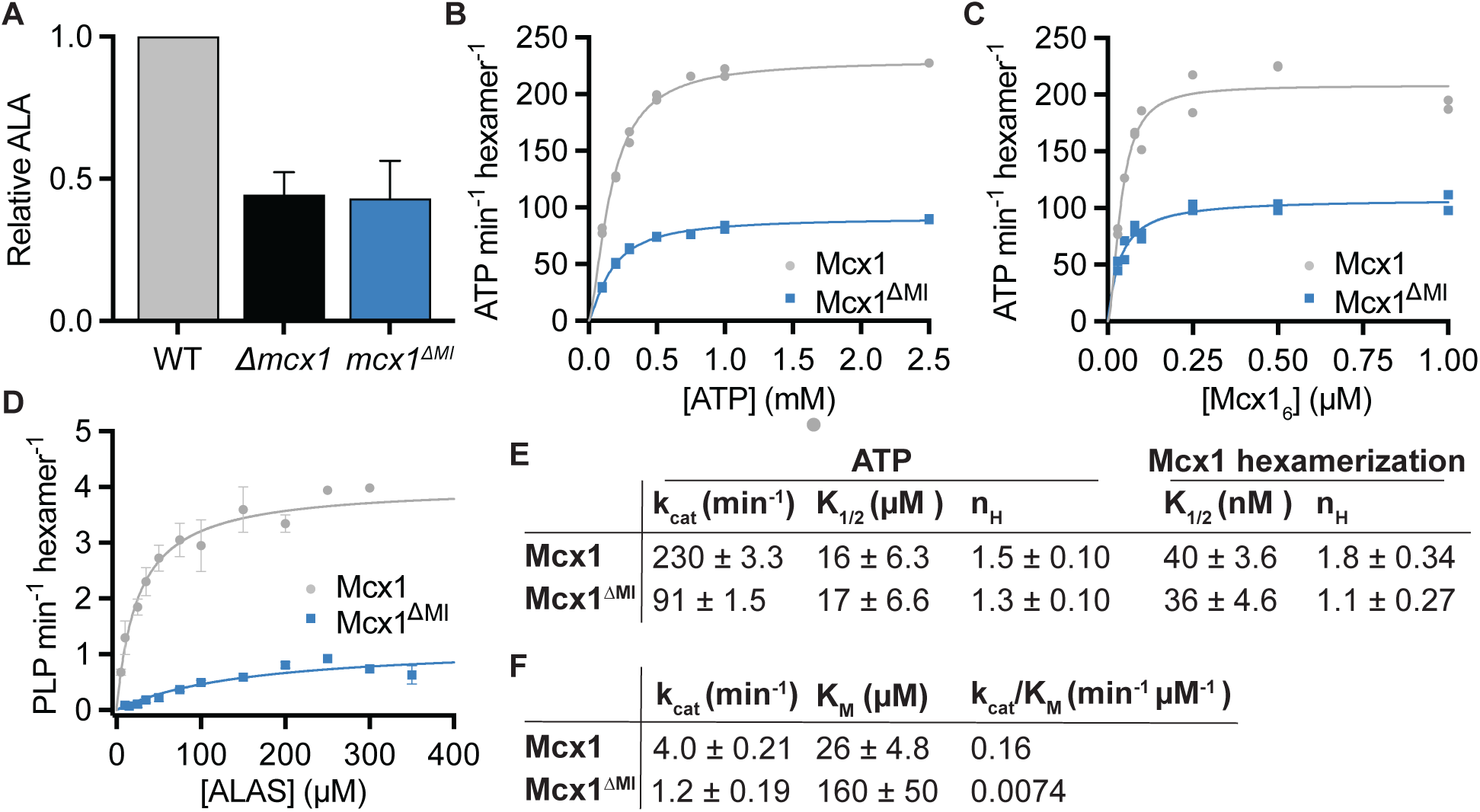
The mitochondrial insertion is critical to support heme biosynthesis through activation of ALAS. (A) Steady state ALA levels in *S. cerevisiae* cell extracts, quantified using modified Ehrlich’s reagent (n = 3, error bars indicate SEM). (B) ATP hydrolysis by 0.5 µM Mcx1 hexamer plotted as a function of ATP concentration and fit with an allosteric-sigmoidal equation (see Methods) (n = 2, all data points displayed). (C) ATP hydrolysis by Mcx1 with 2 mM ATP, plotted as a function of Mcx1 hexamer concentration and fit with an allosteric-sigmoidal equation (n = 2, all data points displayed). (D) Mcx1-stimulated PLP binding to ALAS monitored by fluorescence (ex. 434 nm., em. 515 nm). Data are fit with the Michaelis-Menten equation (n=3, error bars indicate SEM). (E) Kinetic parameters extracted from curve fits in (B-C). (F) Kinetic parameters extracted from curve fits in (D).

Having established that Mcx1 requires the MI to stimulate ALA production *in vivo*, we sought to determine the biochemical mechanism underlying this requirement. We first tested basal, substrate-independent activities of Mcx1: ATP hydrolysis and oligomerization, using recombinantly expressed and purified Mcx1^ΔMI^. We first monitored ATPase activity using an NADH-coupled assay. We found no change in the K_M_ for ATP compared to Mcx1, but observed an approximate 2-fold reduction in k_cat_, indicating that the MI does not contribute to ATP binding, but that its loss reduces the maximal catalytic rate (Fig. 2C). ClpX homologs must hexamerize to form a functional ATPase active site^18,19^; we therefore considered whether the reduced k_cat_ for ATP we observed for Mcx1^ΔMI^ might derive from impaired hexamerization. To assess hexamerization of Mcx1, we monitored ATPase activity across a concentration range of Mcx1 or Mcx1^ΔMI^. We observed equivalent K_1/2_ values for hexamerization of Mcx1 and Mcx1^ΔMI^(Fig. 2D). These data together demonstrate that the MI does not play a role in Mcx1 hexamerization, but supports its maximal ATP hydrolysis rate. Although the MI is not predicted to contact the active site of Mcx1 (Fig. 1B), it might affect the dynamics of the large subdomain of the AAA+ domain to which it connects in a manner that alters catalytic rate.

We next tested how the MI contributes to Mcx1 activation of ALAS *in vitro*. Mcx1 activates ALAS for catalysis by partially unfolding the enzyme, rapidly accelerating the rate of incorporation of the cofactor of ALAS, pyridoxal phosphate (PLP)^16^. We monitored Mcx1-stimulated PLP incorporation into ALAS over a range of ALAS concentrations, detecting ALAS-PLP by the fluorescence of the pyridoxyllysine bond PLP forms at the active site. We observed that truncation of the MI caused a 4-fold reduction in k_cat_ and a 7-fold increase in K_M_ for stimulation of PLP binding by Mcx1, causing a large decrease in catalytic efficiency (Fig. 2D,E). From the substantial increase in K_M_, we infer that the MI is important for recruitment of ALAS. The reduced k_cat_ for ALAS activation could derive from the reduced ATPase activity we observe for Mcx1^ΔMI^, or it could indicate that the MI contributes to how Mcx1 engages ALAS for unfolding.

To isolate features of the MI that are important for its function and determine if the effect of MI truncation on ATPase rate and substrate interactions are separable, we examined its sequence conservation and its predicted structure for coherent features. Although the MI diverges widely in sequence and approximately two-fold in length across eukaryotes, it exhibits some sequence conservation across shorter evolutionary distances (Fig. 1A). Comparing the MI of Mcx1 with other ClpX homologs in ascomycota, two conserved features stand out (Fig. 3A). First, the loop linking two predicted alpha helices is highly conserved, with Phe101 showing nearly complete conservation among ascomycota. Arg84, which maps to the base of the predicted helix-loop-helix (Fig. 3B), also shows broad conservation. To test how these features contribute to Mcx1 ATPase activity and activation of ALAS, we analyzed two Mcx1 variants: Mcx1^loop^, in which all residues in the apical loop (Glu98-Ser105) are mutated to alanine, and Mcx1^R84A^. The maximal ATPase rates of Mcx1^loop^ and Mcx1^R84A^ were equivalent to that of the wildtype enzyme, rather than decreased as for Mcx1^ΔMI^ (Fig. 3C, E). In contrast, both Mcx1^loop^ and Mcx1^R84A^ were impaired in their ability to activate ALAS. Mcx1^loop^ exhibited a four-fold decrease in k_cat_ and five-fold increase in K_M_ for ALAS, similar to the impairment in ALAS activation observed for Mcx1^ΔMI^. Mcx1^R84A^ exhibited a smaller decrease in k_cat_ and increase in K_M_ values for ALAS, causing a three-fold change in the specificity constant (0.16 min^−1^ μM^−1^ for Mcx1; 0.055 min^−1^ μM^−1^ for Mcx1^R84A^) (Fig. 3D, E). These mutations thus uncouple the effects of the MI on ALAS activation from ATPase activity. This uncoupling demonstrates that the requirement for the MI in activation of ALAS is not caused by general impairment of the ClpX ATPase, but instead that the MI contributes to both ALAS recognition (thus affecting the K_M_ for ALAS) and engagement/unfolding by the AAA+ motor (thus affecting k_cat_ for ALAS).

**Figure 3.**
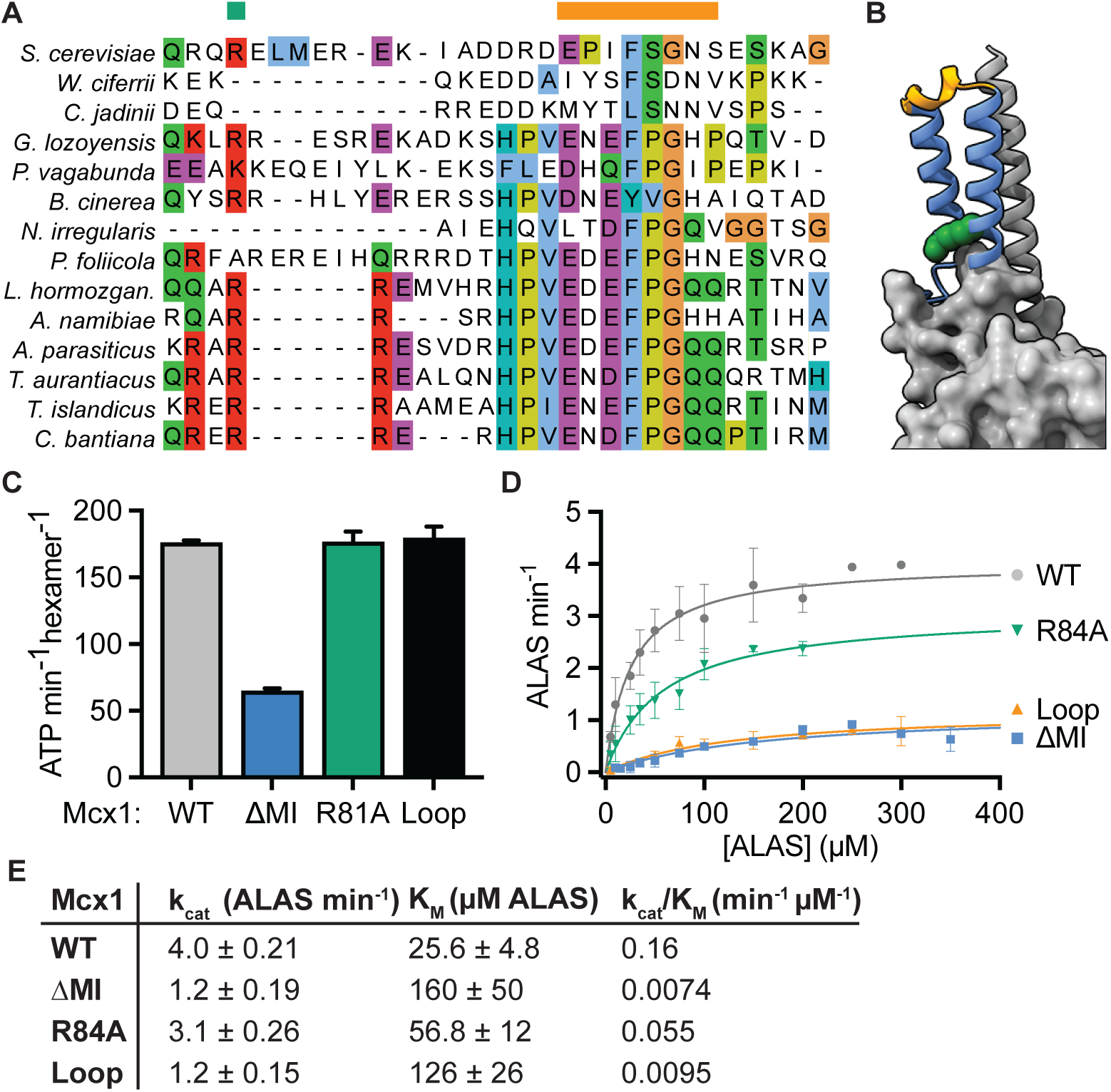
The apical loop of the mitochondrial insertion of Mcx1 mediates both substrate recruitment and engagement for unfolding. (A) A central subsection of the mitochondrial insertion, drawn from a global alignment of ClpX sequences among ascomycota. The green bar indicates the conserved Arg84, and the orange bar indicates the apical MI loop residues. Region displayed corresponds to residues 81-110 in *S. cerevisiae* Mcx1. (B) AlphaFold3 model of Mcx1, in which the MI is displayed in blue ribbon with the apical loop colored orange and Arg84 depicted as green spheres. The N-terminal element is displayed as a gray ribbon and the AAA domain as gray surface. (C) ATP hydrolysis rates of Mcx1 and variants (0.5 µM Mcx1 hexamer, 2 mM ATP, n = 3, error bars indicate SEM). (D) Mcx1-stimulated PLP binding to ALAS, fit with the Michaelis-Menten equation. Parameters extracted from the fit are listed in (E).

We next sought to determine how the MI contributes to the function of human CLPX. The MI emerges from the same location of the AAA+ domain in all mitochondrial ClpX homologs, but its sequence and predicted structure vary considerably between yeast and human homologs (Fig. 1A-C). We prepared a variant of the human CLPX protein analogous to Mcx1^ΔMI^, CLPX^ΔΜΙ^ (with a 3xGly-Ser linker replacing residues 210-280 of CLPX). We first monitored ATP hydrolysis while varying ATP or CLPX concentration to extract kinetic parameters for ATP hydrolysis and CLPX hexamerization. In contrast to the effect of MI truncation on Mcx1 ATP hydrolysis, we observed that the k_cat_ for ATP hydrolysis is mildly stimulated upon MI truncation (Fig. 4A, C). As for Mcx1, human CLPX without the MI exhibited a similar K_M_ for ATP and K_1/2_ for hexamerization (Fig. 4B, C). Therefore, as for the yeast ClpX homolog, we conclude that the MI of human ClpX does not contribute to oligomerization or formation of an ATPase active site. Another group reported that truncation of the CLPX MI, defined as residues 230-304, or mutation of E285 decreased ATPase activity, impaired hexamerization, and nearly abolished degradation of casein by CLPXP^15^. Residues 304 lies well within a strongly conserved region of the AAA+ core domain (ending beyond the region in Fig. 1A), and E285 is within a region of strong structural and moderate sequence conservation. The defects reported from this truncation may thus arise from removal of this more conserved C-terminal region or structural strain imposed by direct fusion of the flanking regions.

**Figure 4.**
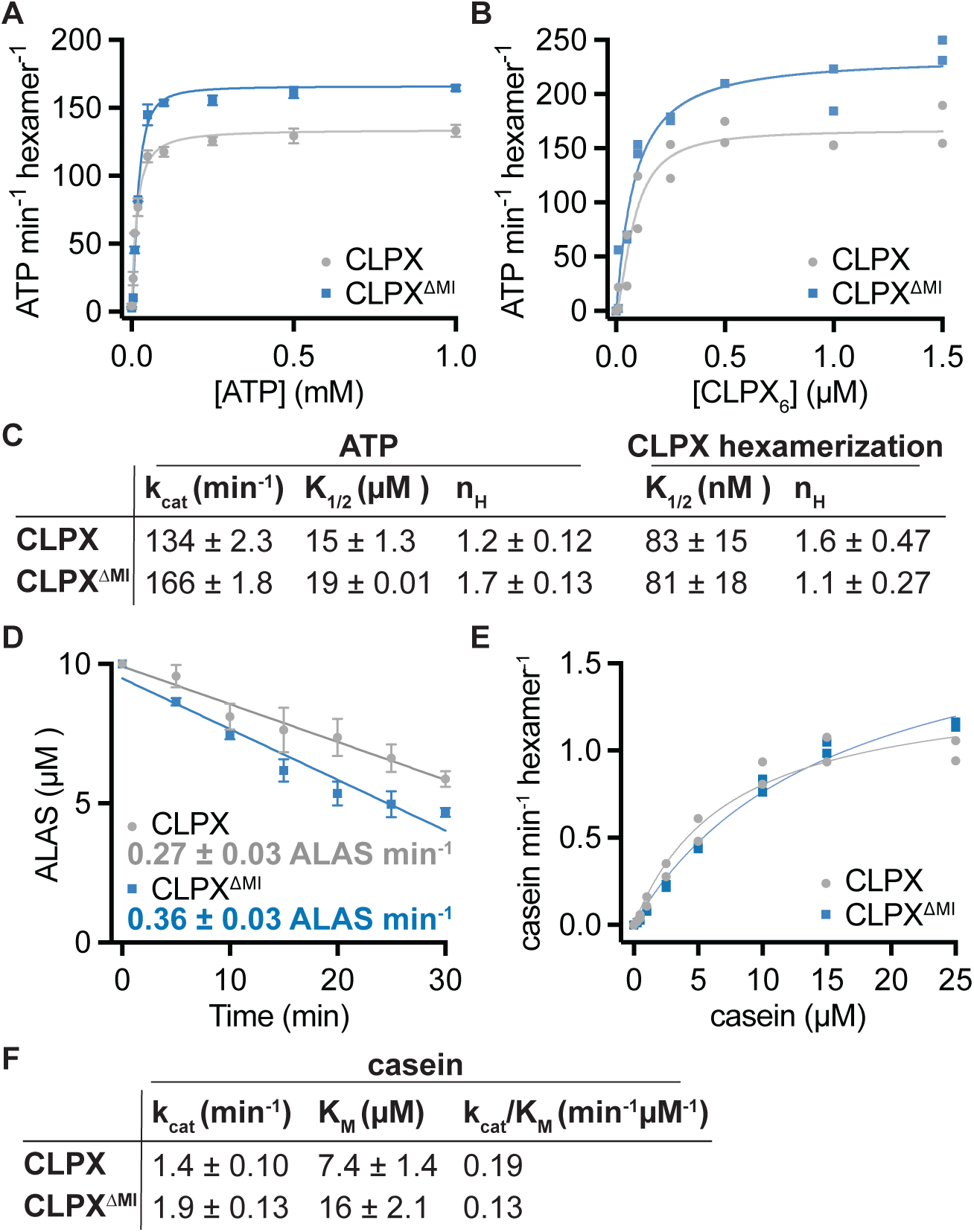
The CLPX MI is not required for degradation of ALAS2, but tunes recognition of casein. (A) ATP hydrolysis by 0.5 µM CLPX hexamer plotted as a function of ATP concentration and fit with an allosteric-sigmoidal equation (n = 3, error bars indicate SEM). (B) ATP hydrolysis by CLPX with 2 mM ATP, plotted as a function of CLPX hexamer concentration, fit with an allosteric-sigmoidal equation (n = 2, all data points shown). (C) Kinetic parameters extracted from curve fits in (A) and (B); error indicates SEM of curve fit. (D) In vitro degradation of 10 μM ALAS2 with 0.5 µM CLPX hexamer, 0.5 µM CLPP 14-mer, 3 µM POLDIP2, 10 µM hemin, and 2 mM ATP. Lines represent linear fits to data, with error bars representing SEM. CLPX-FLAG was used in this experiment to sufficiently distinguish CLPX^ΔMI^ and ALAS2 sizes for analysis by SDS-PAGE. (E) FITC-casein degradation rates by 0.5 µM CLPXP, plotted as a function of FITC-casein concentration and fit with Michaelis-Menten equation (n = 2, all data points shown). (F) Kinetic parameters extracted from curve fits in (D); error indicates SEM.

The moderate increase we observe in the maximal ATP hydrolysis rate of human CLPX after MI truncation was unexpected in comparison with the decreased rate we observed for Mcx1^ΔMI^. One clear difference between CLPX and Mcx1 is that Mcx1 has lost the zinc-finger N-domain that most ClpX homologs, including human CLPX, contain. We considered whether this domain, which projects from the same face of the ClpX hexamer as the MI, might alter the effect of MI truncation. To test this, we also measured the maximal ATPase rate of CLPX lacking the N domain (CLPX^ΔN^) or lacking the N domain and the MI (CLPX^ΔN,MI^). We observed a similar increase in ATPase activity due to MI truncation in CLPX, regardless of the presence of the N-domain (Fig. S2A). The N-domain therefore does not underlie the difference in the effect of MI truncation on the ATPase activity of yeast and human ClpX or otherwise influence the ATPase activity of CLPX. We do observe that AlphaFold3 predicts with moderate confidence that the N-terminal portion of the N-domain in yeast ClpX forms a continuous α helix with the preceding element of the AAA+ domain, whereas the structure of the human MI region is predicted with lower confidence and with a break in the analogous helix between the MI and the AAA+ domain (Fig. 1B-C). These differences in predicted structure additionally suggest that the allosteric communication between the MI and the AAA+ domain is different in these two homologs. In metazoans, mtClpX degrades ALAS in response to heme levels by a negative feedback mechanism, orchestrated by the heme-sensing adaptor protein POLDIP2^14^. We next asked whether the MI contributes to adaptor-mediated feedback degradation of ALAS by mtClpX. We monitored heme-induced degradation of ALAS by CLPX or CLPX^ΔMI^ (with CLPP and POLDIP2). We observed that the rate of degradation was slightly increased upon MI truncation over a ten-fold range of ALAS2 concentration (Fig. 4D, Fig. S2B). The MI therefore is not required for the adaptor-mediated degradation of ALAS2 by CLPXP. The moderate increase in degradation rate could derive from the increased maximal ATPase rate we observe for this variant (Fig. 4A).

We next tested whether the MI contributes to the interaction of POLDIP2 and CLPX. POLDIP2 binds to the N-domain of CLPX^20^, which projects from the CLPX hexamer in close proximity to the MI. AlphaFold3 models of CLPX-POLDIP2 interaction weakly predict a possible additional contact between the MI and POLDIP2 (Fig. S3 A-B). To test this possibility, we determined the affinity of POLDIP2 for CLPX and CLPX^ΔMI^ using fluorescence anisotropy. We observed an equivalent affinity of POLDIP2 for CLPX and CLPX^ΔMI^ (K_D_ of ∼60 nM), indicating that the MI is not required for this interaction (Fig. S3C).

Although no natural substrate of CLPXP other than ALAS2 is well-characterized in vitro, casein is a well-established model substrate of mitochondrial ClpX^21,22^. Casein is the eponymous substrate of the Clp (caseinolytic protease) family of unfoldases, but only some members of this family can recognize it. For example, *E. coli* and *C. crescentus* ClpX homologs are nearly inactive again casein^23,24^. To test if the MI contributes to some aspects of CLPX substrate recruitment, we monitored casein degradation by CLPX or CLPX^ΔMI^ with CLPP. We found that CLPXP^ΔMI^ degraded casein with a two-fold increase in K_M_ and a smaller increase in k_cat_, indicating the MI can contribute to substrate recruitment by CLPX (Fig. 4E, F), but that its loss does not globally impair protein translocation by CLPX as previously proposed^15^.

## DISCUSSION

Here, we have determined that the MI, an element ubiquitously inserted in the AAA+ domain of mitochondrial ClpX homologs, contributes to both substrate recruitment and engagement for unfolding. Although truncation of the MI altered the k_cat_ for ATP hydrolysis (several-fold decrease for the yeast Mcx1, a small increase for human CLPX), formation of an active hexamer of ClpX was unperturbed, as indicated by equivalent K_1/2_ values for ATP and ClpX. Additionally, the effect of the MI on substrate processing can be uncoupled from its effect on ATPase activity: mutation of a region of the MI had a similar effect on substrate processing as a full deletion, but no effect on ATPase activity. We therefore propose that the MI acts primarily to mediate substrate interactions and is not required to assemble a functional ClpX core. The altered ATPase rates we observe in MI-truncated ClpX homologs supports the idea that the MI communicates with the AAA+ motor domain of ClpX. However, because we observe a similar substrate-processing defect in both Mcx1^ΔΜΙ^ and a variant with a mutated MI (Mcx1^loop^) that preserves wildtype levels of ATPase activity, we propose that the primary role of this communication is not to globally regulate ATP hydrolysis, but rather to coordinate substrate engagement with motor activity.

The MI is near-essential for the activation of ALAS by the yeast mtClpX homolog, Mcx1. Deletion or mutation of the MI increases the K_M_ for ALAS by an order of magnitude, but also substantially decreases the k_cat_. Based on the position of the MI on the substrate-recruiting face of ClpX, we had hypothesized a likely contribution to substrate affinity, similar to the previously observed contribution of the bacterial ClpX N-domain and adaptors with multiple bacterial substrates^25^. We unexpectedly observed that the MI is also important for the rate of activation of ALAS by Mcx1. This finding implies that the MI contributes in some way to positioning ALAS for efficient engagement and unfolding by the ClpX motor, or in coordinating the action of the substrate-engaged hexamer.

We find that the mitochondrial insertion more subtly affects several activities of human CLPX. Deletion of the mitochondrial insertion does not affect the affinity of CLPX for POLDIP2, the sole adaptor protein identified for CLPX. POLDIP2-mediated degradation of ALAS as well as degradation of the model substrate casein by CLPXP is slightly accelerated rather than being diminished by MI truncation, perhaps resulting from the slightly increased ATPase rate of CLPX^ΔΜΙ^. We do observe, however, a modest increase in the K_M_ for casein degradation by CLPXP with MI truncation. These data, together with the ubiquitous presence of the MI in all mitochondrial CLPX homologs to our knowledge, suggest that the MI may contribute to substrate recruitment and processing for a subset of CLPX substrates. Because a mechanism of selection for only a single physiological substrate of CLPX, ALAS, has yet been characterized, future investigation of a broader array of candidate CLPX substrates will be necessary to test this idea.

## AUTHOR CONTRIBUTIONS

A.D. and J.R.K. conceptualized this study. A.D., R.B., and J.R.K. designed experiments. A.D. and R.B. performed experiments and analyzed data with oversight from J.R.K. A.D and J.R.K. wrote the manuscript with contributions from R.B.

## ACKNOWLEDGMENTS

We thank Niels Bradshaw, Timo Street, and Joey Davis for valuable discussions and/or critical feedback on the manuscript. This work was supported by National Institutes of Health grant R01GM151332 (J.R.K). A. D. was supported by NIH T32GM007596.

## METHODS

### Yeast strain generation

Yeast genes were modified at chromosomal loci using standard homologous recombination techniques. Strains were grown for experimental purposes in synthetic defined media (SD + CSM, 2% glucose) (Sunrise Science) at 30°C with shaking at 220 RPM.

### ALA measurement

Saturated overnight cultures were diluted to OD=0.1 and grown at 30°C. At approximately OD_600_ 1.0 (range 0.91-1.06), 10 mL x 1 OD equivalent of culture was rapidly filtered, and cell-laden filters were immediately immersed in 0.7 mL ice-cold 7.5% trichloroacetic acid, 13.3 mM N-ethylmaleimide and incubated for 15 minutes. Extracts were withdrawn and centrifuged at 2,000 x g for 10 minutes at 4°C. The supernatant was mixed with 1/3 volume 8% acetylacetone in 2 M sodium acetate and incubated for 90°C for 15 minutes. After cooling for 5 minutes at room temperature, the solution was mixed with an equal volume of modified Ehrlich’s reagent^26^ and incubated at room temperature for 15 minutes. Absorbance at 552 nm and 650 nm was measured; ALA content is proportional to A_552_-A_650_.

### Western blotting

1 mL of cells at OD_600_ = 1.0 (range 0.91-1.06) or equivalent was harvested by centrifugation at 21,000 x g for 30s. The cell pellet was resuspended in 200 μL yeast alkaline lysis buffer (0.1 M NaOH, 50 mM EDTA, 2% SDS, 2% β mercaptoethanol) and incubated for 10 minutes at 90°C. 5 μL 4 M acetic acid was added, the lysate was vortexed for 30s, and then mixed with 50 μL SDS loading buffer (250 mM TRIS-HCl pH 6.8, 50% glycerol, 0.05% bromophenol blue) and incubated for 10 minutes at 90°C. Samples were separated by SDS-PAGE and transferred to PVFD membrane for 1 hour at 100 V. Membrane was blocked with 3% milk in TBST, and probed with either 1 μg/mL anti-FLAG (Sigma, F1804) or anti-Por1 (Thermo Fisher Scientific, 16G9E6BC4). Blots were then probed with 0.5 ng/mL DyLite 800-conjugated goat-anti-mouse antibody, and scanned on a LICOR Odyssey Clx.

### Cloning of vectors for protein purification

Vectors for bacterial expression and purification were generated by cloning the following protein sequences, lacking mitochondrial targeting sequences, and their related variants into pET28b or pET23a with an N-terminal His_6_-SUMO tag, using standard molecular biology procedures: Mcx1 (P38323, 4-520), Hem1 (P09950, 35-548), CLPX (O76031, 65-633), ALAS2 (P22557, 54-587), and POLDIP2 (Q9Y2S7, 52-368). A construct for expressing CLPP (Q16740, 57-277) was previously described^27^.

### Protein expression and purification

Constructs were co-transformed with pRIL into BL21(DE3) *E. coli* for expression. Cultures were grown in Lennox LB at 30°C with shaking at 220 rpm, except for FLAG-tagged CLPX variants, which were grown in Terrific Broth (TB). When the desired OD_600_ was achieved (Mcx1, Hem1, CLPX, ALAS2, CLPP: 0.5, POLDIP2: 0.8), the cultures were brought to the desired temperature for expression (Mcx1, CLPX, and ALAS2: 16°C, CLPP: 37°C, POLDIP2: 18°C), induced with IPTG (Mcx1, CLPX, Hem1, and ALAS2: 1 mM, CLPP: 0.5 mM, POLDIP2, and POLDIP2^V211Acd^: 0.4 mM), and expressed (Mcx1, CLPX, ALAS2, and POLDIP2: overnight; CLPP: 3 hours). For POLDIP2^V211Acd^ expression, acridonylalanine was supplemented to 0.5 mM at time of ITPG induction. After expression, cells were harvested by centrifugation at 5000 x *g* for 10 minutes, washed once in lysis buffer by centrifugation, snap-frozen in liquid nitrogen, and stored at −80°C until purification.

Cell pellets were lysed by two passes through an ice-cold microfluidizer chamber (LM20-30, Microfluidics) in the same base buffer (25 mM HEPES pH 8, 2 mM MgCl_2_, 20 mM Imidazole, 10% glycerol, 1 mM DTT, 0.5 mM PMSF) with varying pressure and salt concentrations (Mcx1, CLPX: 10 kpsi, 100 mM KCl and 400 mM NaCl; Hem1, CLPP, ALAS2: 18 kpsi, 100 mM KCl and 400 mM NaCl; POLDIP2, POLDIP2^V211Acd^: 18 kpsi, 500 mM KCl). Buffers for Hem1 and ALAS2 were additionally supplemented with 20 μM PLP. Buffers for FLAG-tagged CLPX variants were supplemented with 0.01% CHAPS. Proteins were captured with HisPur Ni-NTA resin (Thermo Scientific), washed with additional lysis buffer, and then eluted with lysis buffer supplemented to 250 mM imidazole.

Prior to elution, FLAG-tagged CLPX variants were additionally washed as follows: Resin was washed at room temperature with 50 mL of Ni-NTA lysis buffer with 1 mM DTT, 0.005% CHAPS, and 5 mM ATP, followed with lysis buffer supplemented with 1 mM DTT and 5 mM ATP. A final wash was performed at 4°C with lysis buffer supplemented with 1 mM DTT. The rest of the protein purification proceeded the same as for other ClpX variants above.

Eluted proteins were pooled and incubated with enhanced SUMO protease^28^ during overnight dialysis into their into their respective storage buffers. All proteins were prepared in the same base storage buffer (25 mM HEPES pH 7.6, 10% Glycerol, 1 mM DTT, 2 mM MgCl_2_) with additional salt (Mcx1, CLPX: 300 mM KCl, CLPP: 100 mM KCl, Hem1: 100 mM KCl, ALAS2: 150 mM KCl, POLDIP2: 500 mM KCl). Cleaved SUMO tags were removed by Ni-NTA capture. mtCLPX was additionally purified by flowing through Sepharose Q resin. For PLP-depleted yeast ALAS, 5 mM hydroxylamine was added to SUMO-subtracted protein and incubated overnight at 4°C All proteins were purified by gel filtration (Mcx1, CLPX, CLPP, Hem, ALAS2: Superdex S200, POLDIP2: Superdex S75). Fractions corresponding to desired proteins were concentrated, aliquoted, and snap frozen in liquid nitrogen.

### ATPase assay

Mcx1 or CLPX was mixed with ATP and NADH mix: (20 mM NADH, 150 mM phosphoenolpyruvate, 400 U/mL pyruvate kinase, 400 U/mL lactate dehydrogenase) in a final volume of 25 µL reaction buffer (25 mM HEPES pH 7.4, 150 mM KCl, 5 mM MgCl_2_, 10% glycerol, 1 mM DTT). ATPase activity was monitored by loss of absorbance at 340 nM in a clear, flat-bottom 384-well plate (Corning 3640). To monitor oligomerization, Mcx1 or CLPX concentration was varied and ATP concentration was 2 mM. To monitor kinetic parameters for ATP binding, ATP concentration was varied and measured with 0.5 µM Mcx1 or CLPX hexamer. ATP hydrolysis rates were calculated by dividing the raw absorbance (U/min) by the product of NADH extinction coefficient (6220 M^−1^ cm^−1^), path length (0.328 cm), and enzyme concentration. Experiments were conducted at 30°C for Mcx1, and 37°C for CLPX. Curves were fit with an allosteric-sigmoidal curve where *y* is the ATPase rate, *x* is the concentration of ClpX (hexamerization) or ATP, and H is the Hill coefficient.

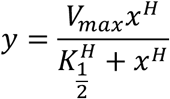

### PLP binding assay

PLP binding to Hem1 was monitored by fluorescence (ex. 434 nm, em. 515 nm) in 25 mM HEPES pH 7.4, 150 mM KCl, 5 mM MgCl2, 10% glycerol with 2 mM ATP, 150 µM PLP, and an ATP regenerating system (5 mM creatine phosphate, 50 µg/mL creatine kinase). Fluorescence was monitored at 30°C in a 384-well plate with 0.5 µM Mcx1 variants and Hem1 concentrations as indicated in individual experiments. Data were fit to the Michaelis-Menten equation to extract k_cat_ and K_M_ paremeters.

### In vitro protein degradation

Degradation of ALAS2 by CLPXP was monitored *in vitro* by incubating ALAS2 at 10 or 1 µM with 0.5 µM CLPX-FLAG (hexamer), 0.5 µM CLPP (14-mer), 3 µM POLDIP2, and 10 µM hemin (Fisher Scientific AAA1116503) in 25 mM HEPES pH 7.6, 150 mM KCl, 5 mM MgCl_2_, 10% glycerol, 1 mM DTT with 2 mM ATP and an ATP regenerating system (5 mM creatine phosphate and 50 mg/mL creatine kinase) at 37°C. Samples were withdrawn at times indicated and immediately mixed with Laemmli SDS sample buffer. Samples were heat-denatured, separated by SDS-PAGE, and stained with SYPRO Red. Gels were imaged on an Amersham Typhoon RGB scanner and quantified in ImageQuant using rolling-ball background subtraction. Rates were extracted using linear fits to all replicates.

To monitor casein degradation, 0.5 µM CLPXP was incubated with indicated concentrations of FITC-casein (Sigma C3777) in the same buffer as for ALAS2 degradation. Degradation was monitored by fluorescence increase (ex. 384 nm, em. 485 nm) in a 384-well plate.

### Fluorescence anisotropy

50 nM POLDIP2^V211Acd^ was incubated with increasing concentrations of CLPX^E359A^ in 25 mM HEPES pH 7.4, 150 mM KCl, 5 mM MgCl2, 10% glycerol, and 1 mM DTT. Measurements were conducted at 37°C in a Fluoromax-4 spectrofluorometer with slit width of 5 nm (ex. 390 nm, em. 444nm). Anisotropy is reported as a function of CLPX_6_ concentration, and fit with a quadradic binding equation where *P* is the polarization value, *a* is the polarization amplitude, *c* is the polarization value without added protein, and *x* is the concentration of CLPX:

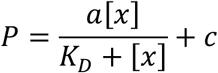

---

**Figure S1.**
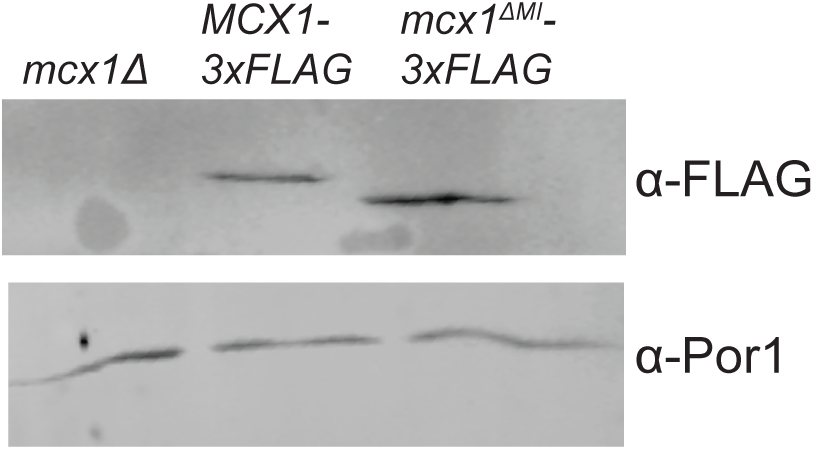
Mcx1-FLAG and Mcx1ΔMI-FLAG are similarly abundant *in vivo*. Western blot of extracts from log-phase *S. cerevisiae*. A C-terminal 3xFLAG tag was appended at the genomic locus of Mcx1. Por1 was probed as a loading control.

**Figure S2.**
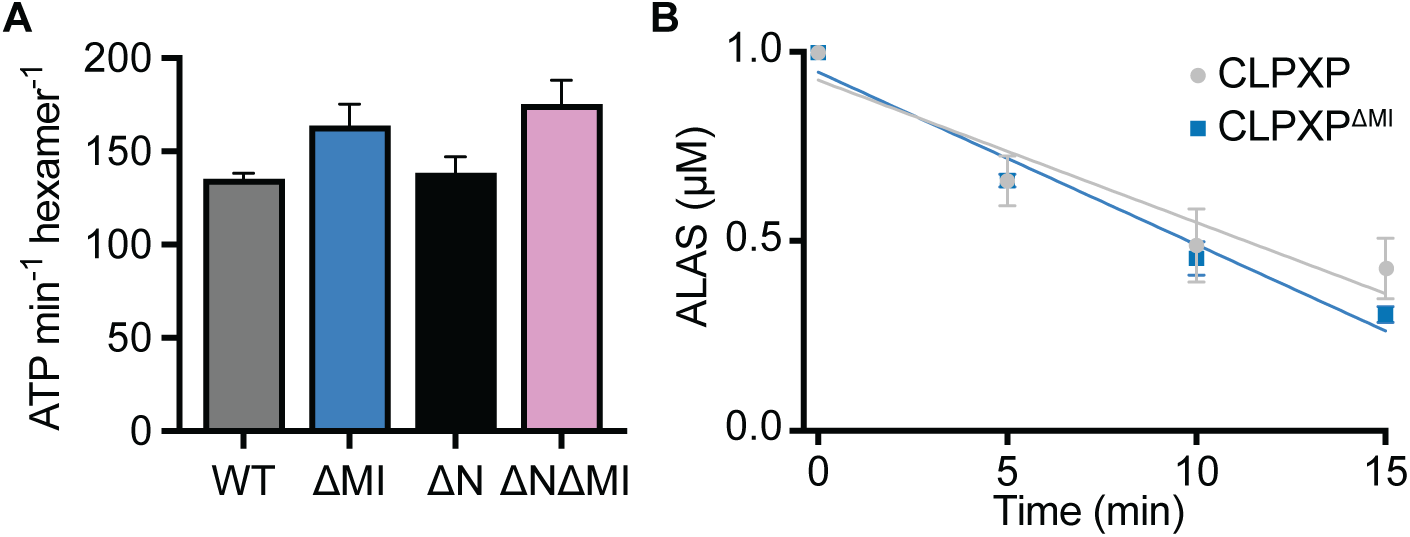
(A) ATP hydrolysis rate of CLPX variants (0.5 µM CLPX hexamer, 2 mM ATP; n = 3, error bars indicate SEM. (B) Degradation rate of 1 μM ALAS2 by CLPX and CLPX^ΔMI^. *In vitro* degradation of 1 μM ALAS2 with 0.5 μM CLPX hexamer, 0.5 μM CLPP 14-mer, 3 μM POLDIP2, 10 μM Hemin, and 2 mM ATP. Linear fits to data are shown; error bars represent SEM for each point (n=3).

**Figure S3.**
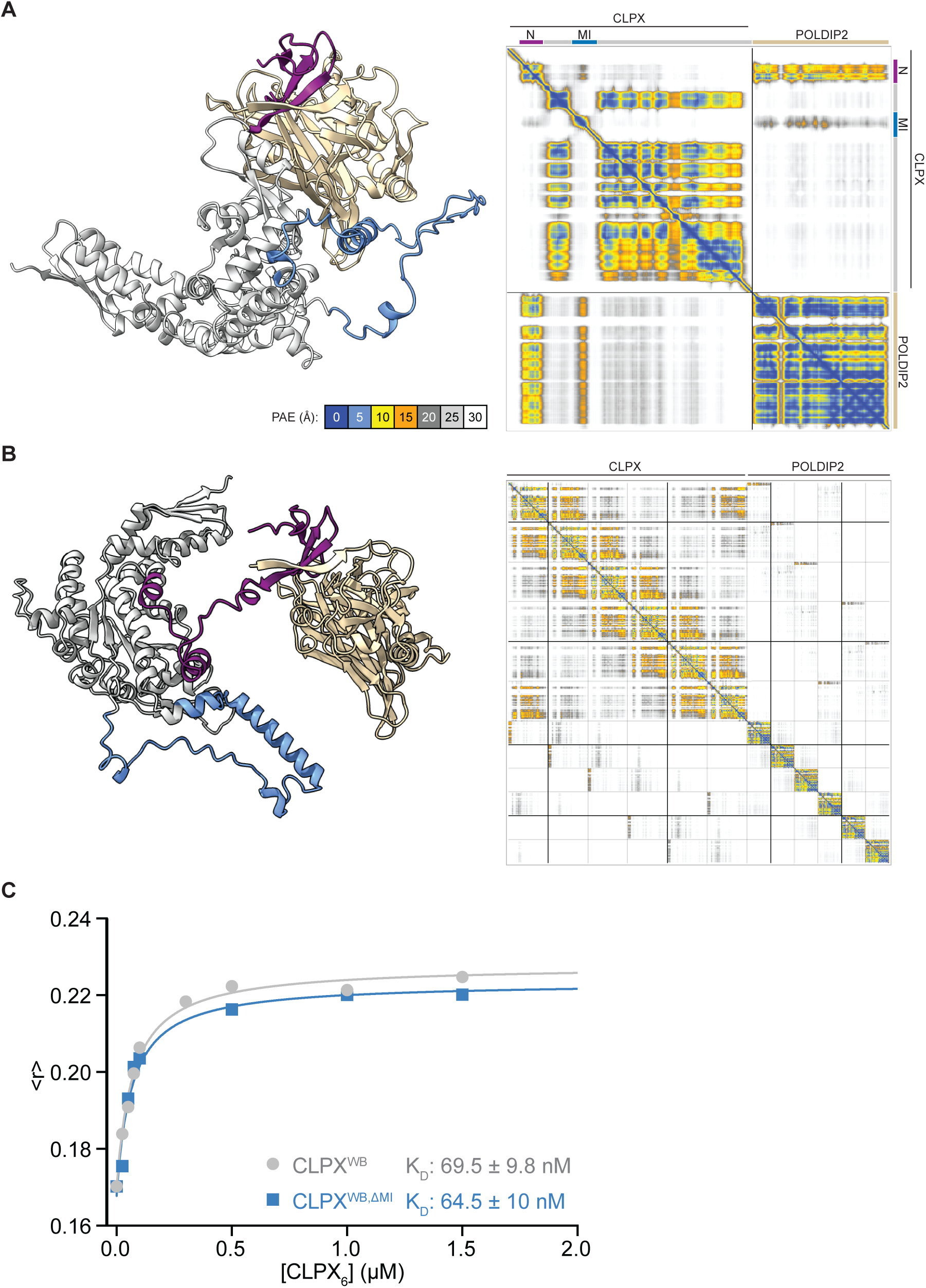
The mitochondrial insertion of CLPX is not required for POLDIP2 binding (A) Left: AlphaFold3 prediction of a CLPX monomer in complex with POLDIP2. CLPX AAA+ domain is colored gray, N-domain is colored purple, and the MI is colored in blue. POLDIP2 is colored orange. Right: PAE map for the structure prediction, with PAE color key at left. (B) Left: AlphaFold3 prediction of CLPX hexamer in complex with six POLDIP2 chains. Right: PAE map for the structure prediction. Coloring is as described in (A). (C) CLPX-POLDIP2 complex formation, monitored by fluorescence anisotropy. CLPX^E359Q^ or CLPX^ΔΜΙ, E359Q^ was titrated against 50 nM POLDIP2^V211Acd^ and fluorescence anisotropy of Acd was monitored (ex. 390 nm, em. 444 nm).

